# Antifungal activity of 2,3-diphenyl-2,3-dihydro-1,3-thiaza-4-ones against *two* human pathogenic fungi

**DOI:** 10.1101/2020.06.27.175711

**Authors:** Livia Liporagi-Lopes, Hany F. Sobhi, Lee J. Silverberg, Radames J.B. Cordero, Arturo Casadevall

**Author notes:** Address correspondence to Livia Liporagi-Lopes.

## Abstract

Invasive fungal diseases are prevalent in immunocompromised individuals in whom current therapies often provide suboptimal results. Additionally, the increased resistance to the available antifungal drugs necessitates a search for new compounds. This study reports the antifungal activity of six 5-, 6-, and 7-membered 2,3-diphenyl-2,3-dihydro-1,3-thiaza-4-ones against *Lomentospora prolificans* and *Cryptococcus neoformans*. Our data showed that some of the compounds tested had a low MIC and damage on the cell surface of the tested fungal species.

## INTRODUCTION

The need for new antifungal compounds reflects the limitations of current therapies, which include frequent therapeutic failures despite prolonged courses and increasing drug resistance [1]. The long-term use of an antifungal drugs can increase the problem of resistance, as already seen with liposomal amphotericin B in an AIDS patient with relapsing/refractory cryptococcosis [2]. Mortality rates for invasive fungal infections remain unacceptably high even when treated with existing drugs. A recent estimate puts the annual death toll from fungal diseases at 1.5 million [3, 4]. The identification of new antifungal compounds is complicated becausee of the similarities in cellular physiology between fungal and animal cells, such that many compounds with antifungal activity are unacceptably toxic to humans [5, 6]. Furthermore, several antifungals have significant interactions with immunosuppressive cells, such as those used in patients after a solid organ transplant (SOT), due to the inhibition of hepatic P450 enzymes [7, 8].

Current therapeutic choices for the treatment of invasive fungal infections are limited to four classes of drugs: 5-fluorocytosine, polyenes including amphotericin B, echinocandins such as anadulafungin, and triazoles [4]. The most recent class of antifungals, the echinocandins, were developd two decades ago [9].

Two fungal pathogens that are very difficult to treat are *Lomentospora prolificans* and *Cryptococcus neoformans*. *L. prolificans* systemic infections are notable for very high morbidity and mortality rates [10]. Cryptococcal meningitis is a global problem resulting in thousands of deaths annually [11]. Poor and late diagnosis, limited access to antifungals, and drug resistance are directly associated with the high fatality rate of cryptococcosis, especially in developing countries [12].

The five, six, and seven-membered 1,3-thiaza-4-ones heterocycles (Fig. 1) are a biologically active group. The most studied are from the the five-membered 1,3-thiazolidin-4-ones group. These compounds are easily prepared and have shown a wide range of activities [29–30]. Derivatives of 1,3-thiazolidin-4-one exhibit antibacterial, antitubercular, anticancer, anti-inflammatory, analgesic, anticonvulsant, antidepressant, antiviral and anti-HIV, trypanocidal (anti-epimastigote), antiarrhythmic, anti-hyperlipidemic, cardiovascular and antidiabetic activities, as well as an agonist of FSH and muscarinic receptors [18,20]. The 2,3-dihydro-1,3-thiazin-4-ones have also displayed antifungal activity [44-49].

**Figure 1.**
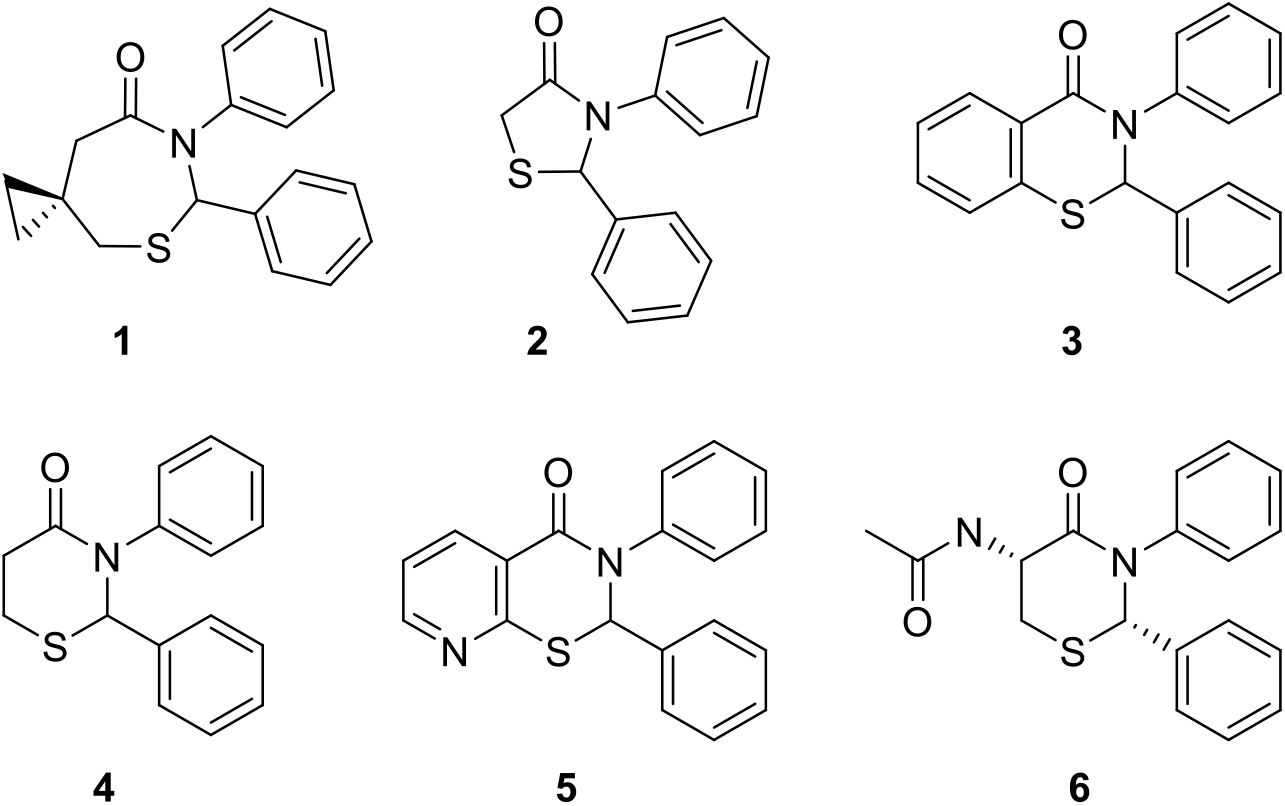
Structures of 2,3-diphenyl-2,3-dihydro-1,3-thiaza-4-ones **1**-**6**.

In this study, we evaluated the potential antifungal activity of six 2,3-diphenyl-2,3-dihydro-1,3-thiaza-4-ones **1-6** (Fig. 1) against conidia of *L. prolificans* and yeasts of *C. neoformans*. The compounds are rings comprised of sulfur at the one position (S1), carbon at two (C2), nitrogen at three (N3), and a carbonyl at four (C4). Each also has benzene rings attached to C2 and N3. Compound **1** is a seven-membered ring that has a spiro cyclopropane at C6. Compound **2** is a five-membered ring. Compounds **3**-**6** are six-membered rings. Compound **3** has a benzene ring fused to C5-C6. Compound **5** similarly has a pyridine ring fused to C5-C6. Compound **6** has an *N*-acetyl moiety attached to C5 and is a single enantiomer as a result of *N*-acetyl-L-cysteine being used in its preparation [21,38]. Compounds **1**-**5** are racemic (Fig. 1).

## METHODS, RESULTS AND DISCUSSION

Stock solutions of the six compounds were prepared at 1000 μg/ml with 100% dimethyl sulfoxide (DMSO, Fisher Scientific Company, USA), followed by serial dilutions to make working antifungal solutions. When combined with the inoculum suspension, the final concentration series ranged from 50 to 0.39 μg/ml. Growth and sterility controls were included for each tested isolate, and *Candida albicans* strain SC5314 was used as a reference-quality control strain in every batch. Finally, the microdilution plates were incubated at 37°C with 180 rpm during 48 (for *C. neoformans* strain H99) or 72 (for *L. prolificans* strain ATCC90853) hours for minimum inhibitory concentration (MIC) determination. The plates were analyzed by measuring the absorbance at 492 nm using a spectrophotometer. Antifungal susceptibility testing was performed to determine the minimal concentration of the compounds necessary to inhibit 50% of the *C. neoformans* and *L. prolificans* growth, according to the Clinical and Laboratory Standards Institute (CLSI) guidelines contained in the M38-A2 document and EUCAST protocol 9.3 [13, 14]. Fluconazole was used as a reference drug. *In vitro* antifungal susceptibility testing is now standardized internationally and the MIC informs the susceptibility or resistance of the organism to the antifungal agent, which can help in treatment decisions [13, 15, 16]. In this context, we tested six 2,3-diphenyl-2,3-dihydro-1,3-thiaza-4-ones against *L. prolificans* and *C. neoformans* and found that thiazepanone **1**, thiazolidone **2**, had better efficacy (lower MIC) against both fungi in comparison with fluconazole. Thiazinone **4** and **5** had a good effect against *C. neoformans*, while benzothiazinone **3** and *N*-acetyl-L-cysteine derived **6** showed no effect (or no effect better than our control, fluconazole) on either fungus (Table 1). Thus, 5-, 6- and 7-membered ring compounds exhibited strong antifungal activity.

**Table 1.** MIC 50 for *C. neoformans* and *L. prolificans* to compounds **1**-**6** and fluconazole. MIC values were obtained from duplicate (compounds **3**-**6**) and triplicate (compounds **1**-**2**) analysis.

|  | MIC 50 (µg/mL) |  |  |  |  |  |  |
| --- | --- | --- | --- | --- | --- | --- | --- |
|  | **1** | **2** | **3** | **4** | **5** | **6** | **Fluconazole** |
| *C. neoformans* | 42.2 | 42.8 | >100 | 47.5 | 27.5 | >100 | 65.0 |
| *L. prolificans* | 31.6 | 28.6 | >100 | >100 | 95.0 | >100 | >100 |

Cytotoxicity for mammalian cells was tested using J774 macrophage-like cells. J774 macrophages cells were plated in 96-well polystyrene tissue-culture plates and incubated for 24 h at 37°C and 5% CO_2_ prior to the addition of compounds **1-6** (final concentration series ranged from 50 to 0.39 μg/ml). After 24 and 48 h incubations, the cell viability was measured by XTT salt method, according to the Assay Guidance Manual [17, 18]. Cytotoxicity assays were done to ascertain whether these compounds could damage J774 macrophages. No significant effect was observed on the macrophage’s viability in the presence of different concentrations of compounds **1**-**6** at 24 and 48h (Fig. 2).

**Figure 2.**
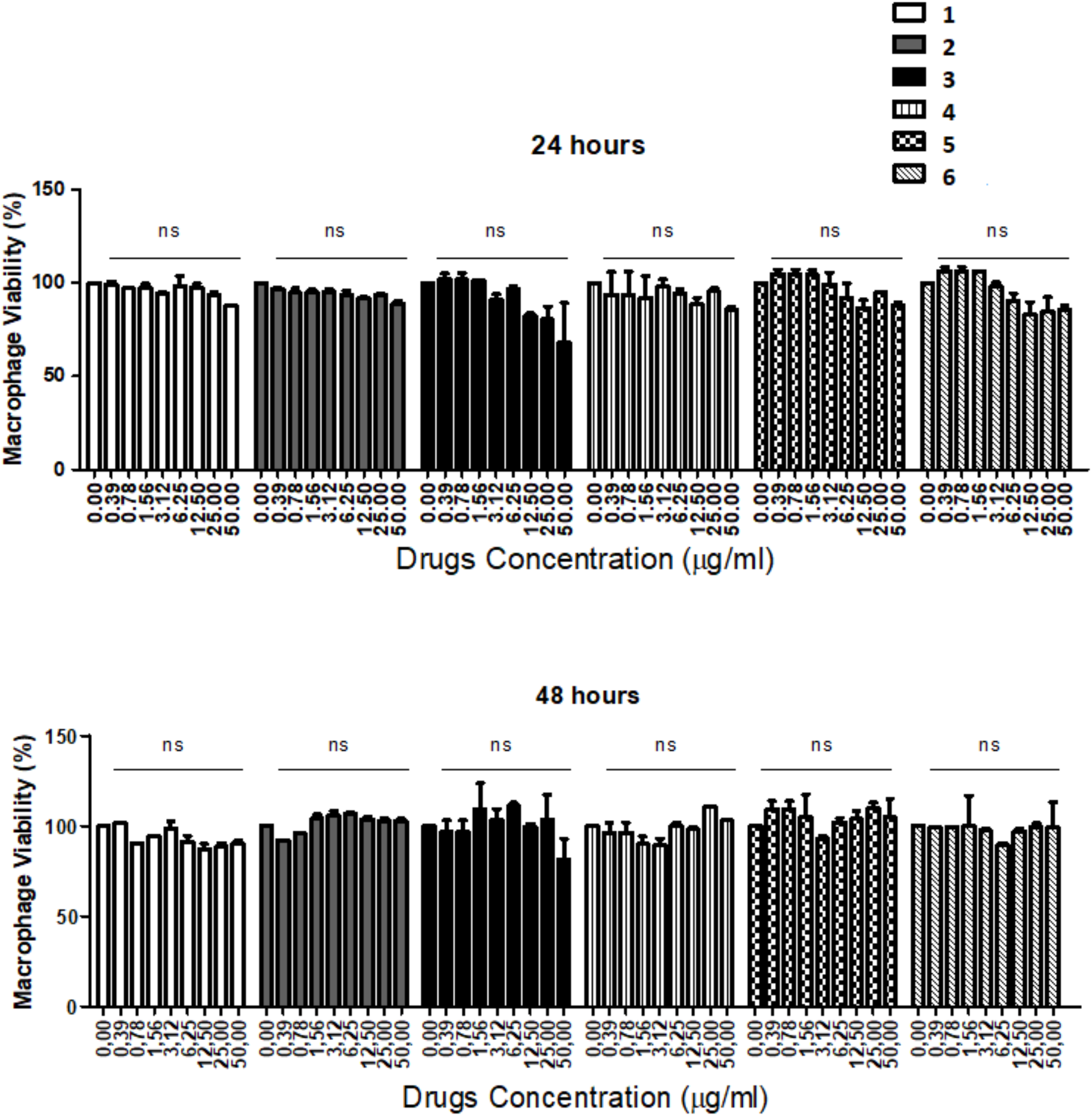
Cytotoxicity assay of macrophages in the presence of compounds **1**-**6** during 24 and 48h, performed according to the Assay Guidance Manual. Statistical analyses were performed using GraphPad Prism version 8.00 for Mac X (GraphPad Software, San Diego CA). One-way analysis of variance using a Kruskal-Wallis nonparametric test was used to compare the differences between groups, and individual comparisons of groups were performed using a Bonferroni post-test. ns: not significant.

Since compounds **1** and **2** were more effective than fluconazole in both fungi, we decided to evaluate the fungal cell surface after cells are incubated with the MIC 50 concentration of compounds **1** and **2** for 48 and 72 h, through SEM [19] and IF [20] techniques. Cell surface damage was found on both *C. neoformans* (Fi. 3, pictures A and C) and *L. prolificans* (Fig. 3, pictures B and D) cells after incubation with compounds **1** and **2**. SEM analysis showed that treated *C. neoformans* cells were often partially collapsed and/or folded (Fig. 3, picture A) while treated *L. prolificans* exhibited the presence of small pores and wrinkles on the cell surface (Fig. 3, picture B). Epifluorescence microscopy of both species using 0.5 mg/mL of Uvitex 2B cell wall staining showed broken cell walls (Fig. 3, pictures C and D). *C. neoformans* yeasts were collapsed and *L. prolificans* cells exhibited pores, suggesting that the inhibitory mechanism of both compounds **1** and **2** may involve interaction with the fungal cell wall.

**Figure 3.**
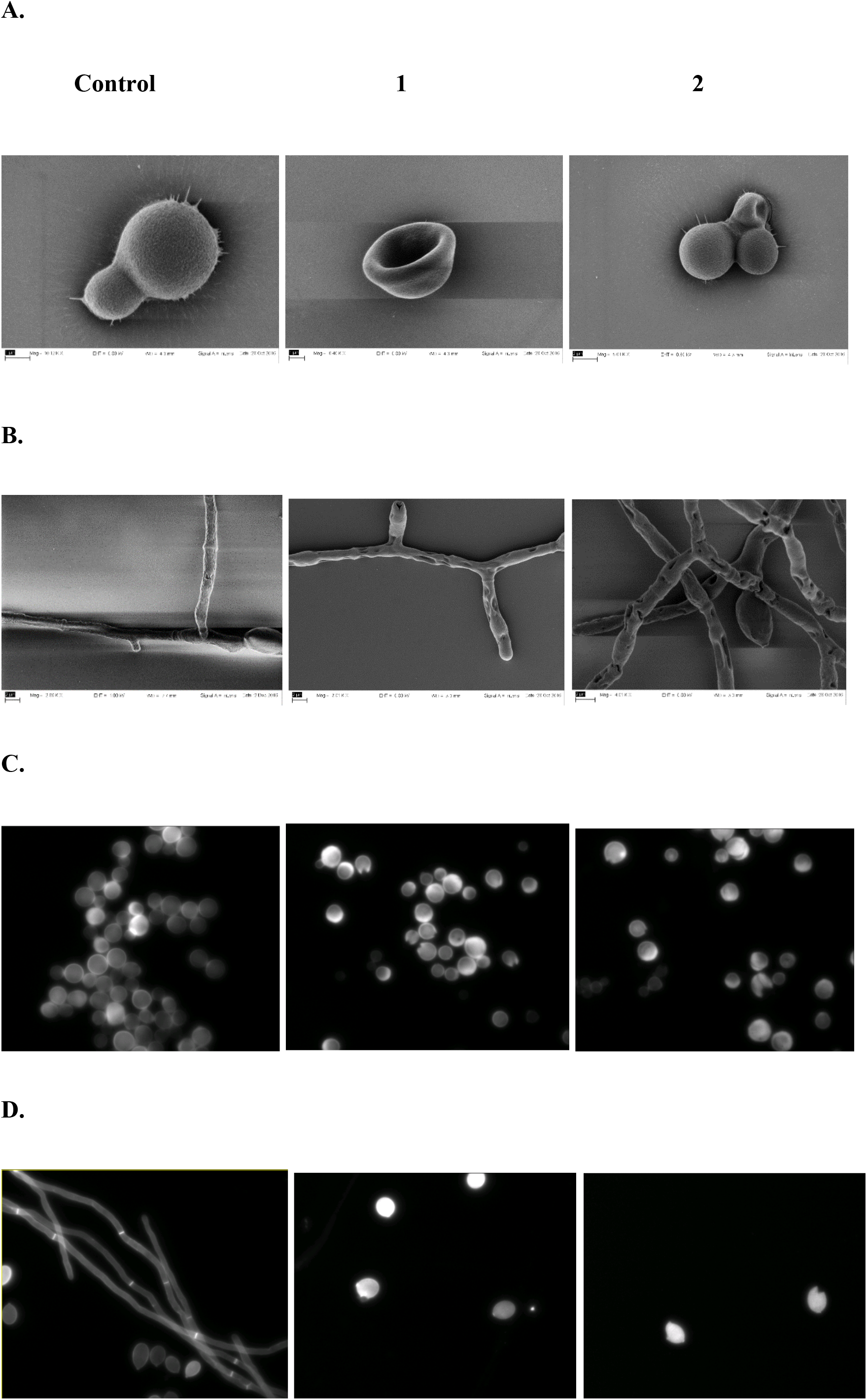
Cell wall damage in *C. neoformans* and *L. prolificans* by compounds **1** and **2** as visualized by SEM and light microscopy. Pictures A and B show SEM of *C. neoformans* and *L. prolificans*, respectively. Pictures C and D show Uvitex 2B staining of fungal cell wall from *C. neoformans* and *L. prolificans*, respectively. Scale bar represents 2 μm.

In summary, our findings suggest that 2,3-diphenyl-2,3-dihydro-1,3-thiaza-4-ones compounds could represent a promising new class for the development of new antifungal therapeutic agents. The mechanism of antifungal action is still under investigation.

## Acknowledgment

We want to thank the Postdoc fellowship from CNPq, Conselho Nacional de Desenvolvimento Científico e Tecnológico – Brasil. Coppin State University, Center for Organic Synthesis, Baltimore, Maryland, USA. Johns Hopkins University, Department of Molecular Microbiology, Bloomberg School of Public Health, Baltimore, Maryland, USA. AC was supported in part by NIH grants AI052733, AI15207 and HL059842.

## References

1. Scorzoni, L., et al., Searching new antifungals: The use of in vitro and in vivo methods for evaluation of natural compounds. J Microbiol Methods, 2016. 123: p. 68–78.

2. de Carvalho Santana, R., et al., Fluconazole Non-susceptible Cryptococcus neoformans, Relapsing/Refractory Cryptococcosis and Long-term Use of Liposomal Amphotericin B in an AIDS Patient. Mycopathologia, 2017.

3. Brown, G.D., et al., Hidden killers: human fungal infections. Sci Transl Med, 2012. 4 (165): p. 165rv13.

4. Campoy, S. and J.L. Adrio, Antifungals. Biochem Pharmacol, 2017. 133: p. 86–96.

5. Ostrosky-Zeichner, L., et al., An insight into the antifungal pipeline: selected new molecules and beyond. Nat Rev Drug Discov, 2010. 9 (9): p. 719–27.

6. Pierce, C.G., et al., Antifungal therapy with an emphasis on biofilms. Curr Opin Pharmacol, 2013. 13 (5): p. 726–30.

7. Dodds-Ashley, E., Management of drug and food interactions with azole antifungal agents in transplant recipients. Pharmacotherapy, 2010. 30 (8): p. 842–54.

8. Denning, D.W., et al., Micafungin (FK463), alone or in combination with other systemic antifungal agents, for the treatment of acute invasive aspergillosis. J Infect, 2006. 53 (5): p. 337–49.

9. Pound, M.W., et al., Overview of treatment options for invasive fungal infections. Med Mycol, 2011. 49 (6): p. 561–80.

10. Cortez, K.J., et al., Infections caused by Scedosporium spp. Clin Microbiol Rev, 2008. 21 (1): p. 157–97.

11. Park, B.J., et al., Estimation of the current global burden of cryptococcal meningitis among persons living with HIV/AIDS. AIDS, 2009. 23 (4): p. 525–30.

12. Rodrigues, M.L., Funding and Innovation in Diseases of Neglected Populations: The Paradox of Cryptococcal Meningitis. PLoS Negl Trop Dis, 2016. 10 (3): p. e0004429.

13. Subcommittee on Antifungal Susceptibility Testing of the, E.E.C.f.A.S.T., EUCAST Technical Note on the method for the determination of broth dilution minimum inhibitory concentrations of antifungal agents for conidia-forming moulds. Clin Microbiol Infect, 2008. 14 (10): p. 982–4.

14. Rollin-Pinheiro, R., et al., Characterization of Scedosporium apiospermum glucosylceramides and their involvement in fungal development and macrophage functions. PLoS One, 2014. 9 (5): p. e98149.

15. Howard, S.J., et al., Frequency and evolution of Azole resistance in Aspergillus fumigatus associated with treatment failure. Emerg Infect Dis, 2009. 15 (7): p. 1068–76.

16. Grossman, N.T. and A. Casadevall, Physiological Differences in Cryptococcus neoformans Strains In Vitro versus In Vivo and Their Effects on Antifungal Susceptibility. Antimicrob Agents Chemother, 2017. 61 (3).

17. Riss, T.L., R.A. Moravec, and A.L. Niles, Cytotoxicity testing: measuring viable cells, dead cells, and detecting mechanism of cell death. Methods Mol Biol, 2011. 740: p. 103–14.

18. Riss, T.L., et al., Cell Viability Assays, in Assay Guidance Manual, G.S. Sittampalam, et al., Editors. 2004: Bethesda (MD).

19. Camacho, E., et al., The structural unit of melanin in the cell wall of the fungal pathogen Cryptococcus neoformans. J Biol Chem, 2019. 294 (27): p. 10471–10489.

20. Fu, M.S., et al., Cryptococcus neoformans urease affects the outcome of intracellular pathogenesis by modulating phagolysosomal pH. PLoS Pathog, 2018. 14 (6): p. e1007144.

